# NeoToxPred: a fine-tuned protein language model with orthologous and length-stratified negative controls for robust toxicity classification

**DOI:** 10.64898/2026.09.18.752542

**Authors:** Seongmin Kim, Min-Seok Kim, Chungoo Park

**Affiliations:** School of Biological Sciences and Technology, Chonnam National University, Yongbong-ro 77, Gwangju 61186, Republic of Korea; Institute of Systems Biology & Life Science Informatics, Chonnam National University, Yongbong-ro 77, Gwangju 61186, Republic of Korea; Dental Science Research Institute, School of Dentistry, Chonnam National University, Yongbong-ro 77, Gwangju 61186, Republic of Korea

**Keywords:** protein toxicity prediction, data curation, protein language model, data leakage, venom proteome

## Abstract

Although protein toxins represent valuable pharmacological templates, predicting toxicity directly from primary sequences is inherently challenging because of their evolutionary dynamics. Active toxins and their benign homologues frequently share identical structural scaffolds and differ by only a few key residue substitutions, making global sequence similarity an unreliable indicator of function. Conventional predictors compound this challenge through flawed negative-set designs that sample non-toxic controls from arbitrary background proteins and reduce redundancy through identity-based clustering under random partitioning. This practice introduces global confounding shortcuts that inflate benchmark performance while failing to capture true functional boundaries. To address these limitations, we developed NeoToxPred, a sequence-only binary classifier that fine-tunes the pretrained ESM-C 600M protein language model end-to-end with a six-layer feed-forward classification head. Although the neural architecture is intentionally standard to ensure computational scalability, the core novelty of the framework lies in its rigorous training-data curation: (i) an orthologous negative set drawn from the same InterPro families under controlled taxonomic proximity to eliminate phylogenetic shortcuts, (ii) a length-stratified negative control set to eliminate sequence length as a predictive cue, and (iii) a strict family-disjoint splitting strategy to prevent performance inflation caused by pattern memorization. On held-out internal benchmarks, NeoToxPred achieved a Matthews correlation coefficient (MCC) of 0.897 and an F1-score of 0.949 on long sequences and an MCC of 0.869 on short peptides, decisively outperforming five recent predictors while maintaining stable performance across varying sequence lengths and evolutionary distances. In a genome-scale external validation using the previously unseen proteome of the redfin waspfish (*Paracentropogon rubripinnis*), NeoToxPred successfully recovered 11 of 16 empirically validated toxins without collapsing into a single-class prediction, whereas structure-dependent baselines identified substantially fewer positives. Computational alanine scanning further confirmed that the framework contextually localizes its predictions to functionally active residues. Taken together, these findings suggest that systematic negative-set design should be prioritized alongside algorithmic complexity in biological sequence classification.

## Introduction

Toxin proteins and peptides constitute a vast yet largely uncharacterized reservoir of bioactive molecules with substantial pharmacological and ecological relevance [1–3]. Because of their high specificity for ion channels, receptors, and distinct physiological pathways, these molecules serve as valuable molecular probes and structural templates for therapeutic development [4,5]. Accurate sequence-based identification of toxicity is also critical for assessing the safety of engineered biologics and screening synthetic sequences for biosecurity risks [6]. In these applications, the continuous influx of candidate sequences, ranging from genome-scale proteomes to synthetic combinatorial libraries and designed variants, far exceeds the throughput of experimental assays. Robust computational frameworks capable of predicting toxicity directly from primary amino acid sequences are therefore essential for prioritizing high-probability candidates for subsequent experimental validation.

However, a defining feature of toxin evolution inherently complicates sequence-based toxicity prediction. Gene duplication is a major mechanism of evolutionary innovation, providing redundant genetic copies that can escape purifying selection and subsequently acquire distinct functions through neofunctionalization or subfunctionalization [7–10]. Animal venoms provide prominent examples of this process, in which a recurring repertoire of physiological protein scaffolds has been independently co-opted for venom delivery across phylogenetically diverse lineages [11,12]. Subsequent duplication events, accompanied by functional divergence and positive selection, have generated large and functionally heterogeneous toxin families [11,13]. In snakes, for instance, venom metalloproteinases expanded from a single, deeply conserved ancestral *ADAM28* gene through tandem duplication and stepwise domain loss [14,15], whereas three-finger toxins arose from a non-secretory squamate Ly6/uPAR ancestor following loss of its membrane-anchoring domain [16]. As these duplicated loci diverge across speciating lineages, the resulting groups of homologous proteins, which share common sequence and structural ancestry and are delineated here as protein families using the InterPro classification framework [17], inevitably encompass both toxic and non-toxic members. Because functional divergence within these protein families is non-uniform and bidirectional [18,19], derived lineages can independently gain or lose specific toxic capabilities [20,21]. Therefore, active toxins and their physiological non-toxic relatives frequently share an identical structural scaffold and differ at only a few critical positions, making similarity to a known toxin an unreliable indicator of toxicity on its own [22,23].

To resolve these subtle sequence-level distinctions, computational toxicity predictors have evolved from early models based on amino acid composition and sequence motifs [24] to deep learning approaches that learn representations directly from sequences, increasingly incorporating structural or graph-based information [25–27]. Several recent methods follow a design similar to that used in the present study, in which a pretrained protein language model is coupled with a neural classifier. For example, ToxDL2.0 incorporates AlphaFold2-derived structural features, whereas CSM-Toxin uses sequence information alone [27,28]. In most cases, the non-toxic class is defined by sampling proteins without toxin annotations or by drawing random or length-matched peptides, after which redundancy is reduced using identity-based clustering tools such as CD-HIT under random partitioning [29]. Because classifier performance depends on the intrinsic difficulty of the examples that must be distinguished, this type of negative-control design can systematically bias evaluation metrics and inflate reported performance [30–32]. This conventional data-curation practice introduces several distinct limitations. First, negative examples obtained from arbitrary background proteomes are often readily separable from true toxins based on broad sequence properties, such as overall amino acid composition, sequence length, or taxonomic origin, rather than on the residue-level determinants of toxicity. Second, identity-based clustering used to reduce redundancy can systematically remove near-neighbor sequences whose phenotypes differ because of only a few key substitutions. This eliminates crucial borderline cases that, although difficult to classify, are essential for defining model decision boundaries.

Furthermore, because threshold-based clustering filters sequences solely according to a fixed similarity threshold, closely related homologous families may still be divided between the training and test sets. Consequently, reported performance may overrepresent memorization of family-specific motifs rather than genuine functional generalization to unseen protein families [33,34]. In parallel, the utility of structure-dependent predictors is often limited by scalability constraints in genome-wide screening, a problem that is particularly relevant for non-model venomous taxa, for which experimentally determined structures are scarce. Taken together, these challenges indicate that the primary bottleneck lies in dataset construction and evaluation design rather than model architecture, underscoring the need for more rigorous benchmarks for protein toxicity prediction.

To address these dataset-level artifacts and better isolate functional decision boundaries, we developed NeoToxPred, a sequence-only binary toxicity classifier that fine-tunes the pretrained 600-million-parameter ESM-C protein language model end-to-end together with a six-layer feed-forward classification head [35]. The neural architecture is deliberately conventional to maintain computational efficiency at scale; the primary contribution of the framework lies in the systematic design of its training data. Negative examples consist primarily of non-toxic proteins from the same protein families as the toxic positives, sampled according to a controlled degree of taxonomic proximity to the host organisms. This design constrains the classifier to rely on fine-grained, toxicity-specific sequence features rather than broad family-level or taxonomic signals. In parallel, a length-matched negative control set removes sequence length as a potential confounding cue. To avoid performance inflation caused by memorization of familiar patterns, we partitioned the dataset using a strict family-disjoint split, ensuring that no protein family appeared in more than one of the training, validation, or test sets. We benchmarked NeoToxPred against five recent predictors on held-out sets of long proteins and short peptides. We further evaluated genome-scale performance using the proteome of the redfin waspfish (*Paracentropogon rubripinnis*), which was excluded from training. Residue-level effects were analyzed by computational alanine scanning, in which changes in predicted toxicity probability following alanine substitution were used to quantify positional importance. Ultimately, this comprehensive evaluation demonstrates that a sequence-only model, when paired with rigorously controlled data, can rival or outperform predictors that rely on complex architectures or multimodal inputs. In doing so, we establish a more stringent and transparent framework for constructing and evaluating future protein toxin classifiers.

## Materials and Methods

### Data preparation

We constructed three sequence pools comprising a positive toxin set, a taxonomically distant orthologous negative set, and a length-matched random negative set. To minimize sequence-homology bias and prevent information leakage across data partitions, all sequences were divided into training, validation, and test sets in a protein family-disjoint manner. The overall dataset construction workflow is schematically illustrated in Fig. 1A. The absolute number of sequences assigned to each split is provided in Supplementary Table S1.

**Figure 1.**
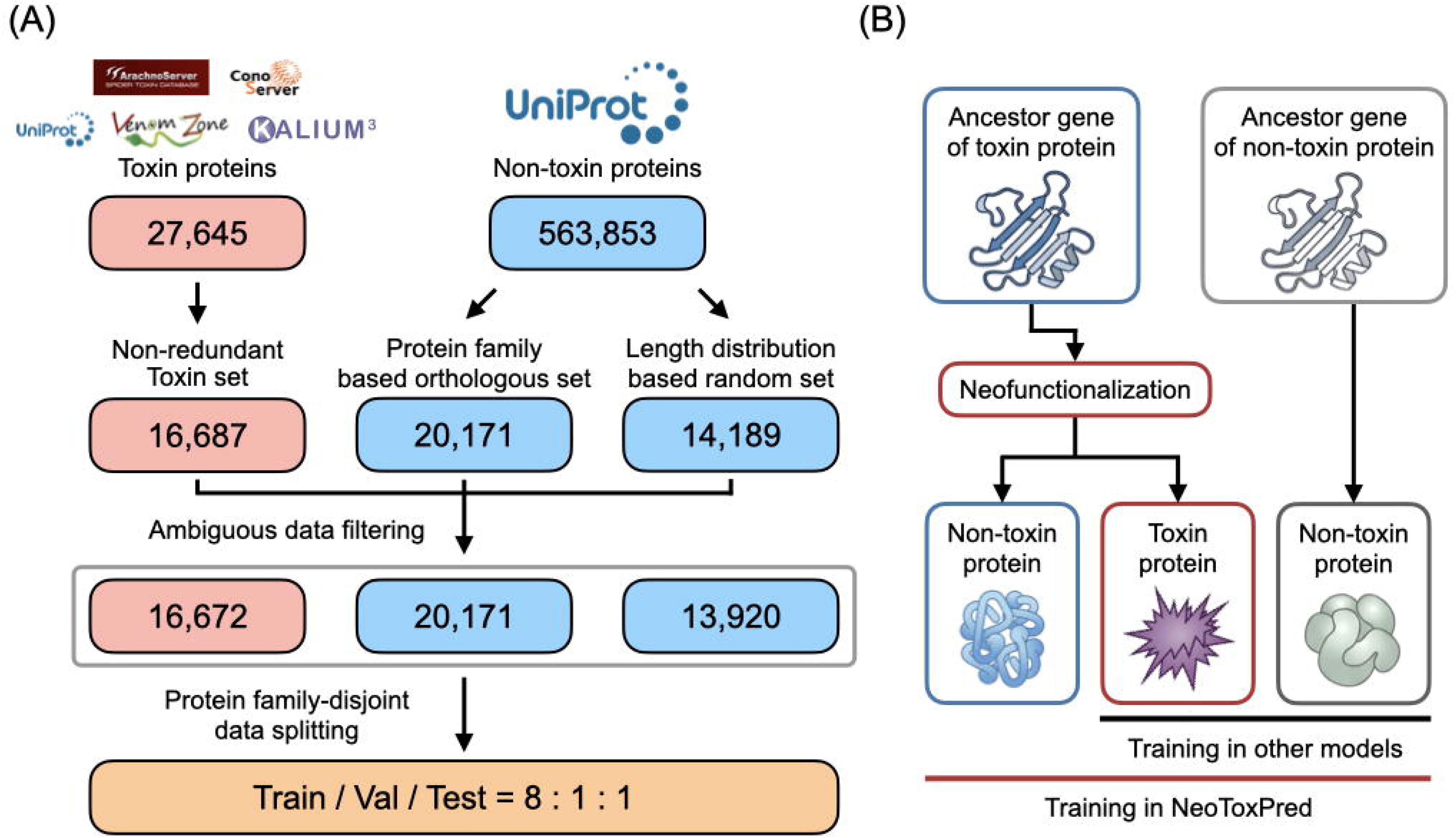
Dataset construction and negative-set design of NeoToxPred. (A) Data curation and partitioning workflow. Toxin sequences compiled from five source databases and non-toxin sequences retrieved from UniProt/Swiss-Prot were curated into a non-redundant positive set, a protein-family-matched orthologous negative control set, and a length-stratified negative control set. Following removal of ambiguous entries, strict family-disjoint partitioning generated the training, validation, and test datasets at an 8:1:1 ratio. Numbers indicate the sequence counts remaining at each curation step. (B) Evolutionary relationships among the sequence classes evaluated during training. Gene duplication followed by neofunctionalization can generate a toxin protein and a non-toxic homologue that descend from a common ancestral gene and retain a shared structural scaffold. Non-toxin proteins of unrelated ancestry share neither this evolutionary relationship nor the corresponding sequence similarity. Horizontal lines indicate the sequence classes contrasted during training: conventional models pair toxins with unrelated background proteins (black), whereas NeoToxPred pairs toxins with both same-family non-toxic homologues (orthologous negative set) and unrelated background proteins (length-stratified negative set) (red).

### Positive (toxin) set

The positive dataset was compiled from five primary sources: ArachnoServer [36], ConoServer [37], UniProtKB/Swiss-Prot [38], VenomZone [39], and KALIUM [40], which contributed 1,838, 8,526, 7,871, 7,794, and 1,616 sequences, respectively. The UniProt subset was retrieved using the query “(((cc_tissue_specificity:venom) OR (keyword:KW-0800)) AND (reviewed:true)).” All databases were accessed in June 2025 (UniProt release 2025_06). After duplicate entries across sources were removed, the resulting non-redundant set comprised 16,687 sequences. Subsequent filtering to remove entries with ambiguous data (see below) yielded a final set of 16,672 toxin sequences.

### Negative-set rationale

The composition of the negative set can substantially influence the performance of toxicity classifiers, and benchmark estimates may be inflated when negative examples are trivially distinguishable from positives based solely on sequence length, amino acid composition, or evolutionary origin [31]. This issue is particularly consequential for toxins because gene duplication followed by neofunctionalization can place toxins and their non-toxic homologues within the same protein family, where they share a common ancestral gene and structural scaffold (Fig. 1B). Standard identity-based redundancy reduction using CD-HIT [41] does not resolve these issues because it removes sequences symmetrically from the positive and negative sets while discarding closely related toxin–non-toxin pairs and leaving homology between training and held-out data unaddressed. Accordingly, CD-HIT filtering was not applied. To control separately for sequence-length and evolutionary biases, we designed two complementary negative sets: an “orthologous negative set,” which targets evolutionary bias by selecting non-toxin sequences that are evolutionarily comparable to toxins, and a “length-matched random negative set,” which targets sequence-length bias by matching the length distribution of the toxin set. Both negative sets were derived from an initial pool of 563,853 candidate non-toxin proteins retrieved from UniProtKB/Swiss-Prot using the query “NOT ((cc_tissue_specificity:venom) OR (keyword:KW-0800) OR (keyword:KW-0020)) AND (reviewed:true).”

### Orthologous negative set

Non-toxin sequences were selected from the same InterPro family or homologous superfamily as the toxin proteins, as identified using InterProScan (v5.68-100.0) [17]. To minimize the risk of including unannotated or cryptic toxic activities, we applied a strict taxonomic exclusion criterion under which orthologous sequences were drawn only from organisms belonging to clades phylogenetically distant from each toxin-producing host. Orthologous candidate sets were constructed at four taxonomic stringency levels (kingdom, phylum, class, and order) by excluding organisms that belonged to the same group as the toxin-producing host at the corresponding level, yielding 826, 5,137, 10,188, and 20,171 sequences, respectively. Taxonomic identifiers were verified by BLASTp (2.15.0+) [42] searches against the NCBI NR database (Fig. S1). The order-level set (20,171 sequences), which provided the largest pool while maintaining interclade separation from venom-producing taxa, was used as the primary orthologous negative set for model training, whereas the kingdom-, phylum-, and class-level sets were retained for downstream analyses of taxonomic robustness.

### Length-matched random negative set

As an independent control for sequence-length bias, we additionally constructed a length-matched random negative set. Toxin sequences were first grouped into 5-amino-acid length bins, and non-toxin sequences were then randomly sampled from the remaining UniProtKB pool, stratified to reproduce the per-bin length distribution of the positive set (Fig. S2). The resulting random negative set comprised 14,189 sequences. Within each negative set, duplicate sequences were removed independently.

### Ambiguous-label filtering and final set sizes

To ensure unambiguous class assignments, we first identified sequences that appeared in both the positive and negative pools and removed them from all sets. After this filtering step, the final dataset comprised 16,672 toxin sequences, 20,171 orthologous negative sequences, and 13,920 length-matched random negative sequences.

### Family-disjoint splitting

The dataset was partitioned at the InterPro family level such that no family was shared between the training, validation, and test splits. Families were allocated to the three splits in an 8:1:1 ratio (training:validation:test), subject to the constraint that each InterPro family appeared in only one split. Using different random seeds, we generated five independent family partitions and derived the corresponding sequence-level splits for each. These five seeded replicates were used to quantify variability throughout the subsequent analyses.

### External validation set

For genome-scale external validation, we used the proteome of *Paracentropogon rubripinnis* (redfin waspfish), a venomous tetrarogid scorpaeniform fish whose dorsal-fin spines contain toxin-secreting glands [43]. The draft genome assembly, protein-coding gene set (36,228 genes), and catalog of 26 dorsal-spine-specific toxin candidates identified from an eight-tissue transcriptome were obtained from a recently reported genomic resource for this species (Lee SG, Park C. Submitted; NCBI BioProject PRJNA1122111, GenBank JBEHBK000000000.1). The toxin class was derived from these 19 candidates by retaining 16 sequences that contained only the 20 standard amino acids represented in the positive training set and had lengths between 150 and 1,600 amino acids. The non-toxin class was constructed from the remaining *P. rubripinnis* proteins by applying the same InterPro family and length filters and excluding sequences with non-canonical or ambiguous residues, yielding 196 non-toxin sequences.

### The architecture of NeoToxPred

NeoToxPred consists of a pretrained protein language model backbone followed by a feed-forward neural network (FFNN) classification head (Fig. 2). The model performs binary toxicity classification directly from amino acid sequence input. Given an input amino acid sequence *s* = (*a*_1_, *a*_2_,…, *a_L_*) of length *L,* we used the ESM-C 600M model [35] to obtain contextualized residue-level embeddings:

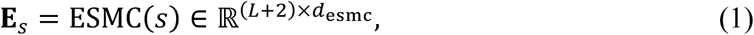

where *d*_esmc_ = 1,152 and the additional two positions correspond to the special <cls> and <eos> tokens. We extracted the <cls> token embedding **x** = **E***_s_*[0, :] ∈ ℝ^1152^as a fixed-length representation of the entire sequence. The ESM-C backbone parameters were jointly fine-tuned with the downstream classifier (i.e., freeze_backbone = False). The CLS embedding **x** was passed through a six-layer FFNN. The first layer projected the input from 1,152 to 4,096 dimensions, after which the hidden dimensions progressively decreased to 2,048, 1,024, 512, 256, and 128. Each layer consisted of a linear projection, a Gaussian Error Linear Unit (GELU) activation [44], and layer normalization:

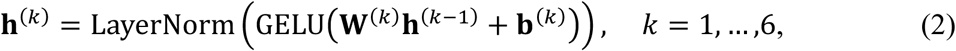

with **h**^(0)^ = **x**. The final hidden representation **h**^(6)^ ∈ ℝ^128^ was projected to a scalar logit through a linear classification head:

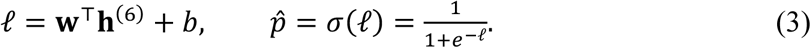

A class label was assigned by thresholding *p̂* at 0.5.

**Figure 2.**
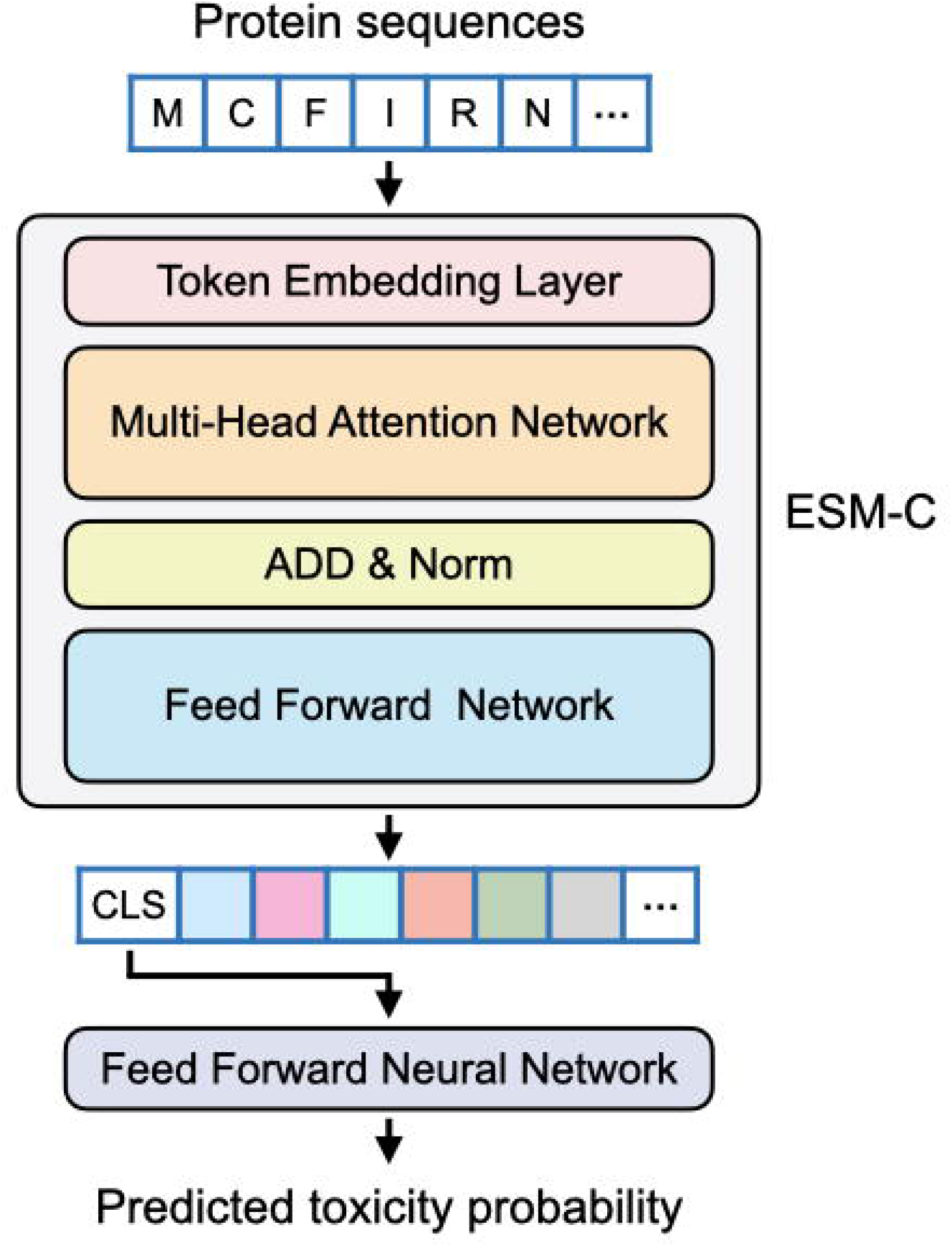
Schematic architecture of the NeoToxPred framework. An input amino acid sequence is encoded using the ESM-C protein language model. The extracted [CLS] token embedding is subsequently fed into a feed-forward neural network (FFNN) to yield the final predicted toxicity probability.

### Training procedure

The model was trained by minimizing the binary cross-entropy loss computed directly from raw logits using BCEWithLogitsLoss, which combines the sigmoid activation and cross-entropy computation in a numerically stable formulation.

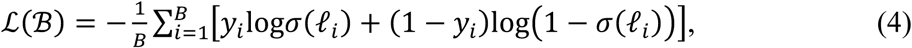

where *l_i_* denotes the raw logit for sequence *s_i_* and *σ*(•) is the sigmoid function. Parameters were optimized using the Adam optimizer [45] with a fixed learning rate (*α*) of 1 × 10⁻^4^, *β*_1_= 0.9, *β*_2_= 0.999, and weight decay (*λ*) of 0. Training was conducted in single precision (FP32) on a single NVIDIA H100 (80 GB) GPU. To prevent overfitting, early stopping was applied: training was terminated when the validation loss failed to improve for five consecutive epochs (patience = 5), up to a maximum of 30 epochs. The model weights from the epoch with the lowest validation loss were retained as the final parameters for each replicate. One model was trained on each of the five family-disjoint splits, yielding five independent replicates. Unless stated otherwise, all metrics reported in this work are presented as the mean ± standard deviation across the five replicates.

### Evaluation metrics

We evaluated model performance using six confusion-matrix-derived metrics. Let TP, FP, TN, and FN denote true positives, false positives, true negatives, and false negatives, respectively, at a decision threshold of 0.5. Recall (sensitivity), specificity, precision (positive predictive value), and negative predictive value (NPV) were defined as:

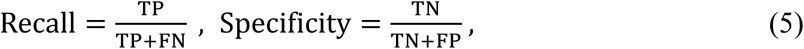

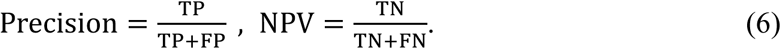

The F1 score, defined as the harmonic mean of precision and recall, was computed as:

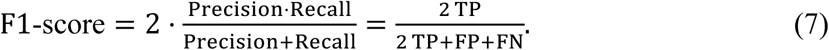

The Matthews correlation coefficient (MCC), which incorporates all four cells of the confusion matrix and is particularly appropriate for class-imbalanced binary classification, was computed as:

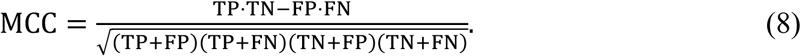

### Baseline models

NeoToxPred was benchmarked against five recently published toxicity predictors evaluated on identical test and external validation sets: CSM-Toxin [28], ToxinPred3 [24], ToxiPep [25], ToxDL 2.0 [27], and tAMPer [26]. Two additional methods, ATSE [46] and ToxIBTL [47], were excluded from the comparison because of limited accessibility. Where method-specific constraints applied, the evaluation was adjusted accordingly. ToxiPep, which accepts only sequences of 50 residues or fewer, was evaluated on the corresponding length-filtered subset. For structure-based baselines, three-dimensional structures were predicted using ColabFold (v1.5.5) [48], and the top-ranked model according to pLDDT score was used as input. All baselines were run using the authors’ published checkpoints and default hyperparameters; no baseline model was retrained on NeoToxPred’s training data.

Performance was assessed under two evaluation regimes. For the internal evaluation, we drew two subsets from the held-out test set: a 4,665-sequence subset composed exclusively of the 20 standard amino acids and a 2,356-sequence subset of standard-amino-acid sequences shorter than 50 residues, providing a peptide-scale evaluation. For the external evaluation, we used the *P. rubripinnis* set described in the External validation set section (16 toxins and 196 non-toxins). Reported metrics for every model-evaluation pair were MCC, F1 score, precision, recall, specificity, and NPV.

### Computational alanine scanning

To quantify the contribution of individual residues to predicted toxicity, we performed computational alanine scanning on two representative sequences from the test set: cone-snail conopressin/conophysin isoform 3 (UniProt ID: A0A4Y5X1A7) and human oxytocin-neurophysin 1 (UniProt ID: P01176). Both sequences belong to InterPro families IPR000981 (neurohypophysial hormone) and IPR036387 (neurohypophysial hormone domain superfamily); A0A4Y5X1A7 is annotated as venom-derived, whereas P01176 is not. Pairwise alignment of the two sequences was performed using Clustal Omega (v1.2.4) [49]. For each residue in the sequence, we defined a position-resolved perturbation score:

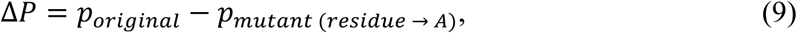

where *p_ori_*_I*inal*_ denotes the toxicity probability assigned by NeoToxPred and *p_mutant_* _(*residue* →_ *_A_*_)_ denotes the toxicity probability assigned to the sequence in which the residue was substituted with alanine. A positive *ΔP* therefore indicates a decrease in predicted toxicity upon substitution, whereas a negative *ΔP* indicates an increase. Positions at which the wild-type residue was already alanine were assigned *ΔP* = 0.

## Results

### Performance is robust across evolutionary distances of the orthologous negative set

A central design choice in NeoToxPred is the use of an orthologous negative set, in which non-toxin sequences are drawn from the same InterPro family or homologous superfamily as the positive toxin sequences. This design forces the model to learn features that distinguish toxins from their closest non-toxic relatives rather than superficial signals that separate venom-derived proteins from arbitrary background sequences. To evaluate whether model performance depends on the phylogenetic composition of the negative set, we constructed four orthologous negative sets with progressively decreasing evolutionary distances between the negative sequences and the toxin-producing hosts. Each set excluded non-toxin sequences at one of four taxonomic ranks (kingdom, phylum, class, or order), as described in the Materials and Methods. The order-level set was the most stringent of the four, retaining only orthologs from organisms that were phylogenetically closest to the venom-producing taxa.

NeoToxPred maintained robust performance across all four taxonomic stringency levels, with an MCC ≥ 0.820 and an F1 score ≥ 0.906. All six confusion-matrix-derived metrics remained within narrow ranges (Supplementary Table S2). Performance was highest at the order level, reaching an MCC of 0.873 ± 0.013 and an F1 score of 0.934 ± 0.012. The order-level set was also the largest (*n* = 20,171); although the larger training set may itself have contributed to performance, model accuracy did not decline at any of the four levels as the phylogenetic gap between toxins and negatives narrowed. Because a model exploiting taxonomic shortcuts would be expected to perform less well under these conditions, the consistency across levels suggests that NeoToxPred does not rely strongly on such features. Based on this robustness, the order-level set was adopted as the primary orthologous negative set for all subsequent analyses.

### Benchmarking against state-of-the-art predictors on long-sequence and short-peptide test sets

NeoToxPred was benchmarked against five recently published toxicity predictors using two subsets of the held-out test set. The compared models were ToxDL2.0, CSM-Toxin, ToxinPred3 (Hybrid), tAMPer, and ToxiPep. The long-sequence subset (*n* = 4,665) contained sequences composed exclusively of the 20 standard amino acids, whereas the short-peptide subset (*n* = 2,356) was restricted to sequences shorter than 50 residues. ToxiPep, which accepts only sequences of 50 residues or fewer, was evaluated on the short-peptide subset only. CSM-Toxin and ToxDL2.0 additionally require predicted three-dimensional structures, which were generated using ColabFold (Materials and Methods).

On the long-sequence subset, NeoToxPred achieved the highest values for five of the six evaluation metrics (MCC, F1 score, precision, specificity, and NPV), reaching an MCC of 0.897 and an F1 score of 0.949 (Fig. 3A). ToxDL2.0 ranked second with an MCC of 0.775 and comparable recall (0.953 vs. 0.947), while showing lower values for the remaining metrics. The other predictors, ToxinPred3 (Hybrid), tAMPer, and CSM-Toxin, showed substantially lower MCC and F1 scores on this subset (Fig. 3A). On the short-peptide subset, NeoToxPred again achieved the highest MCC (0.869) and F1 score (0.937) (Fig. 3B).

ToxDL2.0 yielded marginally higher recall (0.972) and NPV (0.954), but at the cost of lower precision and specificity, indicating a tendency toward overprediction for short peptides. A direct comparison of the two subsets revealed a model-dependent length effect. All baselines except tAMPer, which was trained exclusively on sequences shorter than 100 residues, showed reduced MCC on the short-peptide subset relative to the long-sequence subset (Fig. 3A, B). NeoToxPred’s MCC declined by only 0.028 between the two subsets, from 0.897 on the long-sequence subset to 0.869 on the short-peptide subset, representing the smallest reduction among all evaluated models. This stability across sequence lengths is consistent with the use of a length-stratified random negative set during training (Supplementary Fig. S2), which reduced the opportunity for the model to associate short sequence length with a non-toxic label.

**Figure 3.**
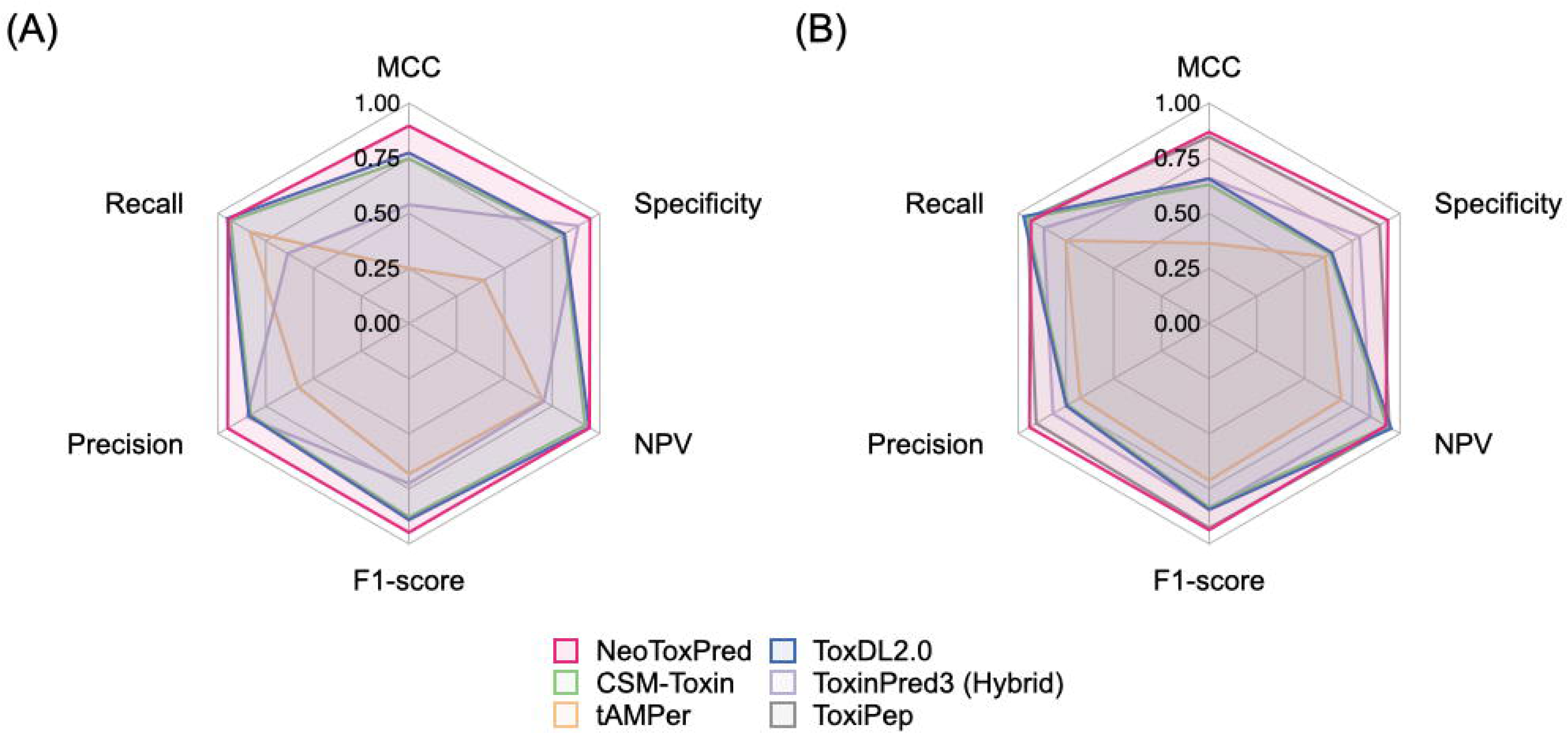
Performance benchmarking of NeoToxPred against five baseline predictors on held-out test datasets. Radar plots show six classification metrics (MCC, F1 score, precision, recall, specificity, and NPV) evaluated on (A) the long-sequence subset (*n* = 4,665) and (B) the short-peptide subset (sequences ≤ 50 residues; *n* = 2,356). ToxiPep was evaluated exclusively on the short-peptide subset.

### External validation on the *Paracentropogon rubripinnis* venom proteome

To assess generalization beyond the held-out in-domain test set, we tested NeoToxPred and four baseline models (ToxDL2.0, CSM-Toxin, ToxinPred3 (Hybrid), and tAMPer) on the proteome of *P. rubripinnis*, a venomous tetrarogid scorpaeniform fish whose draft genome and multi-tissue transcriptome have recently been reported (Lee SG, Park C, Submitted). The external validation set was constructed as described in the Materials and Methods. The toxin class comprised 16 proteins with sequence homology to known animal toxins and dorsal-spine-enriched expression, whereas the non-toxin class comprised 196 proteins from the same proteome that shared InterPro family membership with curated toxins but lacked dorsal-spine expression enrichment. ToxiPep was excluded because the sequence lengths in this dataset exceeded its 50-residue input limit.

All models exhibited lower absolute performance on this external set than on the in-domain test set (Fig. 4A). NeoToxPred achieved the highest MCC (0.480) and F1 score (0.512). tAMPer reached the highest recall (1.000) but the lowest specificity (0.020), reflecting near-complete classification of all sequences as toxins. ToxinPred3 (Hybrid) showed the opposite pattern, with a specificity of 0.969 but no recovered toxins (recall = 0.000). ToxDL2.0 and CSM-Toxin yielded recall values of 0.438 and 0.188, respectively, with specificities comparable to that of NeoToxPred (0.918). The confusion-matrix panel was restricted to NeoToxPred, ToxDL2.0, and CSM-Toxin because tAMPer and ToxinPred3 (Hybrid) produced single-class predictions (Fig. 4B). Among the three retained models, the numbers of correctly classified non-toxin sequences were similar: 180, 189, and 180 for NeoToxPred, ToxDL2.0, and CSM-Toxin, respectively. The numbers of correctly classified toxin sequences differed more substantially, with NeoToxPred identifying 11 of 16, compared with 7 for ToxDL2.0 and 3 for CSM-Toxin.

**Figure 4.**
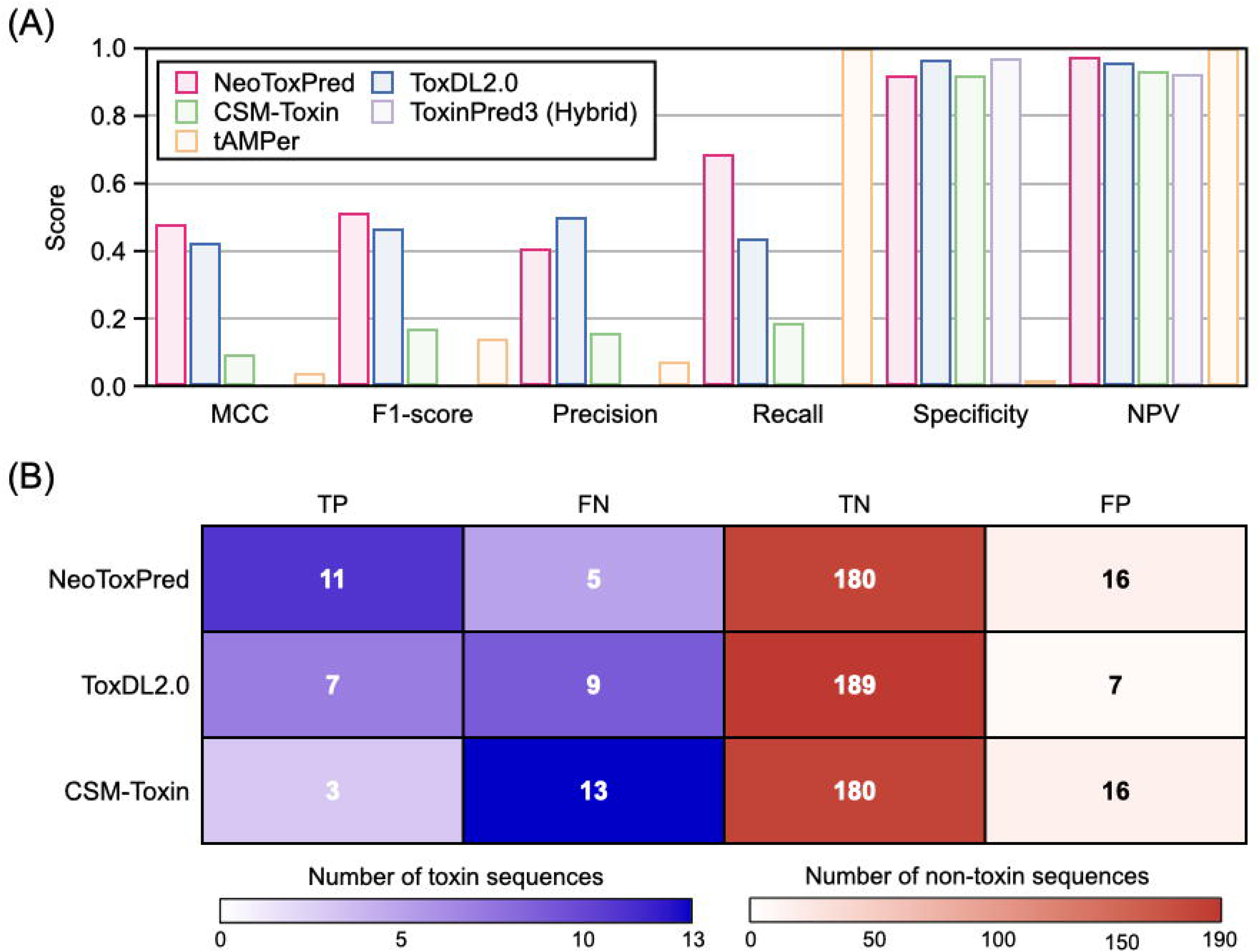
External validation on the *Paracentropogon rubripinnis* proteome (16 toxin candidates and 196 non-toxin sequences). (A) Comparative evaluation of six classification metrics (MCC, F1 score, precision, recall, specificity, and NPV) for NeoToxPred and four alternative predictors. (B) Confusion matrices showing true positives (TP), false negatives (FN), true negatives (TN), and false positives (FP) for NeoToxPred, ToxDL2.0, and CSM-Toxin. tAMPer and ToxinPred3 (Hybrid) are omitted because they produced near-single-class predictions.

### Residue-level perturbation as an illustrative case

To illustrate how NeoToxPred distinguishes a toxin from a non-toxic homolog at the residue level, we selected a representative pair from the same InterPro family in the held-out test set: the venom-derived conopressin/conophysin isoform 3 (UniProt A0A4Y5X1A7) and human oxytocin-neurophysin 1 (UniProt P01176), which share 45.74% pairwise sequence identity (Fig. 5A). Each precursor contains a mature nonapeptide within a longer prepropeptide sequence (CFIRNCPKG in A0A4Y5X1A7 and CYIQNCPLG in P01176) that constitutes the bioactive form of the molecule. NeoToxPred correctly classified both precursors, assigning baseline toxicity probabilities of 0.992 to A0A4Y5X1A7 and 0.007 to P01176. We then applied computational alanine scanning along the full length of each sequence and calculated the position-resolved perturbation score *ΔP* (Materials and Methods, Eq. 9).

**Figure 5.**
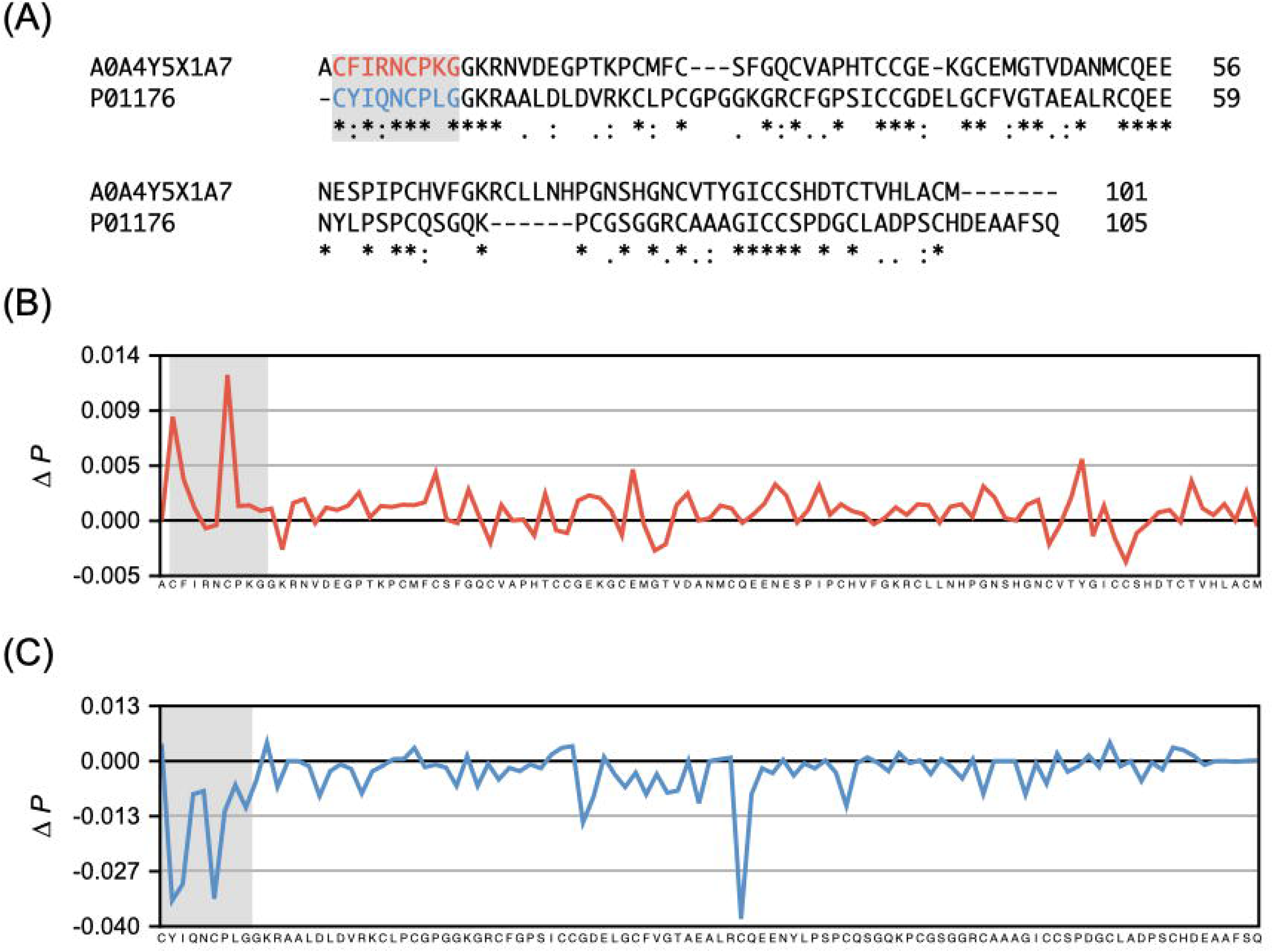
Sequence-level interpretability of NeoToxPred through computational alanine scanning. (A) Pairwise sequence alignment of conopressin/conophysin isoform 3 (UniProt ID: A0A4Y5X1A7, venom-derived) and oxytocin-neurophysin 1 (UniProt ID: P01176, non-toxic), with the mature nonapeptide region highlighted in gray. (B, C) Position-resolved perturbation scores (ΔP) quantifying changes in predicted toxicity probability following alanine substitution at individual residues along the precursor sequences of (B) A0A4Y5X1A7 and (C) P01176.

Within the aligned mature nonapeptide region (highlighted in Fig. 5A), the two precursors shared six of nine residues (positions 1, 3, 5, 6, 7, and 9; C-I-N-C-P-G) and differed at three positions (2, 4, and 8). The corresponding *ΔP* profiles in this region differed markedly between the two sequences. For A0A4Y5X1A7, *ΔP* was strongly positive across the nonapeptide, with the largest contributions at the cysteine and phenylalanine positions (*ΔP* = +0.009 and +0.003, respectively; Fig. 5B), whereas *ΔP* values elsewhere in the precursor remained close to zero. For P01176, the corresponding nonapeptide showed predominantly negative *ΔP* values, indicating that the wild-type residues at these positions supported the non-toxin prediction (Fig. 5C). At the six positions where the two sequences shared identical residues, NeoToxPred nevertheless assigned *ΔP* values with opposite signs. In addition, a second region of strongly negative *ΔP* (−0.038) was observed in P01176 within the LRCQEE segment downstream of the nonapeptide (Fig. 5C), with no comparable peak at the aligned position in A0A4Y5X1A7.

## Discussion

NeoToxPred achieved the highest MCC and F1 score in every comparison reported here: an MCC of 0.897 on the long-sequence test set, 0.869 on the short-peptide subset, and 0.480 on the external *P. rubripinnis* proteome. This advantage persisted under increasingly stringent evaluation conditions (Figs. 3 and 4). These findings suggest that factors beyond architectural differences contribute substantially to the observed performance gap. In particular, the advantage is unlikely to be attributable primarily to model architecture. ToxDL2.0, the closest-performing baseline (MCC = 0.775 on the long-sequence subset), uses the same general strategy as NeoToxPred, namely, a pretrained protein language model followed by a neural classifier, while additionally incorporating two inputs not used by NeoToxPred: a graph neural network operating on AlphaFold2-predicted structures and protein-domain embeddings. Thus, a sequence-only model outperformed a structure-aware model within the same broad modeling paradigm, suggesting that the performance difference is not simply explained by architectural complexity or input modality. More broadly, studies in machine learning have shown that, among models of similar classes trained for the same task, the design and quality of the training data can have a greater influence on performance than the specific model architecture [30,32]. In biological sequence classification, the composition of the negative set can likewise have a substantial effect on benchmark performance [31]. We therefore propose that the construction of the training data is a major factor underlying the performance of NeoToxPred. Standard identity-based redundancy reduction with CD-HIT [29] removes sequences above a fixed pairwise-identity threshold, but it does not control which non-toxin proteins remain close to toxins in sequence space and therefore does not directly determine the biological difficulty of the classification boundary. When negatives are drawn from arbitrary non-toxin proteins, a classifier may distinguish the two classes using broad compositional or phylogenetic signals rather than the sequence differences that separate a toxin from a closely related non-toxin protein. The two negative sets used here were designed to reduce these potential shortcuts. The orthologous negative set places the classification boundary between toxins and non-toxins from the same protein families and comparable taxonomic groups, whereas the length-matched random set reduces the association between sequence length and class label. In addition, partitioning the training, validation, and test sets so that no protein family is shared among them prevents the model from resolving these biases simply by memorizing family-level patterns. Together, these design choices make the classification task more biologically realistic and encourage the model to rely on sequence features that distinguish toxins from closely related non-toxins rather than on readily exploitable dataset artifacts.

NeoToxPred performed consistently across the four orthologous negative sets, with an MCC ≥ 0.820 and an F1 score ≥ 0.906 at every taxonomic level (Table S2). The four sets differed in the evolutionary distance between the negative sequences and the toxin-producing hosts. At the kingdom level, the negatives were derived from more distantly related organisms, whereas at the order level they were derived from the most closely related taxa, resulting in greater sequence similarity between toxins and non-toxins. The order-level set therefore represents the most challenging of the four conditions because the model has less opportunity to exploit broad evolutionary differences and must instead distinguish toxins from closely related non-toxin proteins. Performance was highest at this level, with an MCC of 0.873 ± 0.013 and an F1 score of 0.934 ± 0.012, while remaining stable across the other taxonomic levels. This consistency suggests that the model can distinguish the two classes without relying predominantly on broad evolutionary origin. The finding is particularly relevant to toxin discovery because proteins of practical interest often include non-toxic homologues of known toxins rather than unrelated background proteins, and models trained primarily against arbitrary negatives may be less informative for this type of discrimination. The order-level set was also the largest (*n* = 20,171 versus 826 at the kingdom level), and the greater number and diversity of training examples may have contributed to its higher performance. Nevertheless, the stability of performance across all four taxonomic levels indicates that the model was not dependent solely on evolutionary distance as a classification cue. A similar pattern was observed with respect to sequence length. NeoToxPred showed one of the smallest performance differences between the long-sequence and short-peptide subsets, with an MCC decrease of only 0.028, from 0.897 to 0.869 (Fig. 3). The other models, except ToxinPred3 (Hybrid) and tAMPer, showed lower MCC values on the short-peptide subset.

Such length dependence can arise when the negative sequences used during training are not matched to the length distribution of the toxin sequences. If short proteins are more prevalent among the negatives than among the positives, a classifier may learn an association between short length and the non-toxin class, which can reduce its ability to recognize short toxins during testing. ToxinPred3 (Hybrid) and tAMPer were exceptions among the baselines, likely in part because they were trained on sequences shorter than 35 and 100 residues, respectively, and therefore operated under different length distributions. ToxDL2.0 illustrates a related consequence of uncontrolled length distributions: it achieved higher recall on short peptides (0.972) but lower precision and specificity, a pattern consistent with greater assignment of the toxin label to short sequences. The relatively stable performance of NeoToxPred across sequence lengths is consistent with the use of a length-matched negative set (Fig. S2) and may reflect the influence of this training-data design rather than an intrinsic advantage in processing short sequences. This distinguishes NeoToxPred from length-restricted tools such as tAMPer and ToxiPep.

Evaluation against the *P. rubripinnis* proteome provided a more stringent test of generalization beyond the in-domain test set. This species was completely absent from the training dataset, and the toxin candidates were defined using dorsal-spine expression rather than curated toxin annotations. Although all models showed lower performance on this challenging external dataset than on the held-out test set (Fig. 4), their different failure patterns provide additional insight into their practical behavior. Two baseline models produced near-single-class predictions: tAMPer classified nearly every sequence as a toxin (recall = 1.000, specificity = 0.020), whereas ToxinPred3 (Hybrid) showed the opposite tendency, recovering no toxin sequences (recall = 0.000) while achieving high specificity. Such highly imbalanced predictions substantially limit a model’s usefulness for genome-scale screening because they provide little discrimination among candidate proteins. The practical value of toxin screening lies in prioritizing a manageable set of candidates for downstream experimental validation, which requires meaningful separation between likely toxins and non-toxins. Among the models that retained predictions for both classes, NeoToxPred, ToxDL2.0, and CSM-Toxin achieved similar numbers of correctly classified non-toxin sequences but differed substantially in toxin recovery. NeoToxPred identified 11 of the 16 putative toxins, compared with 7 for ToxDL2.0 and 3 for CSM-Toxin. Consequently, NeoToxPred achieved the highest MCC (0.480) among the evaluated models. These results indicate that NeoToxPred can recover a substantial fraction of toxin candidates while maintaining specificity sufficiently high to produce a more useful candidate set for downstream validation.

Aggregate metrics establish that NeoToxPred can discriminate toxins from closely related non-toxins, but they do not reveal which sequence features contribute to individual predictions. To examine this question at single-residue resolution, we applied computational alanine scanning. In experimental mutagenesis, truncating a side chain beyond the β-carbon can help infer its contribution to molecular function [50,51]. Analogously, this in silico perturbation approach systematically replaces each residue with alanine and quantifies the resulting change in the predicted toxicity probability (*ΔP*). This framework is widely used to interpret sequence-based deep learning models, in which changes in model output following localized sequence perturbations can provide information about positional importance [52,53]. We applied this approach to a pair of homologous proteins from the same InterPro family in the held-out test set: the venom-derived conopressin/conophysin isoform 3 (A0A4Y5X1A7) and the non-toxic human homologue oxytocin-neurophysin 1 (P01176), which share 45.74% sequence identity (Fig. 5A). Both precursors contain a mature nonapeptide domain, and NeoToxPred correctly classified both proteins. In the venom-derived precursor, alanine substitutions produced a localized change in *ΔP* that was concentrated almost exclusively within the mature nonapeptide region (Fig. 5B), suggesting that this region contributes substantially to the model’s classification of the sequence.

Comparative analysis further showed that identical substitutions at conserved positions in the two mature nonapeptides produced opposite changes in *ΔP*, attenuating the predicted toxicity in one protein while increasing it in the other (Fig. 5B, C). If the model relied solely on the identity of individual residues, identical substitutions would be expected to produce more uniform directional effects. Instead, the divergent responses indicate that NeoToxPred evaluates residues in their sequence context, consistent with the non-local sequence dependencies captured by protein language models such as ESM-C [54]. These residue-level observations complement the aggregate performance metrics by illustrating how the model can distinguish homologous proteins with different predicted toxicity through sequence-context-dependent effects rather than global sequence similarity alone. However, computational alanine scanning is an interpretability approach rather than a direct measurement of biological function. Because each *ΔP* value reflects an isolated single-residue substitution, the analysis cannot resolve epistatic interactions among residues [55,56], even though the underlying transformer architecture can model sequence dependencies jointly. Accordingly, *ΔP* should be interpreted as a measure of the model’s response to sequence perturbation rather than as an empirical biophysical property. The identified positions therefore represent hypotheses that can be prioritized for subsequent experimental mutagenesis.

Beyond the analysis-specific limitations described above, the study as a whole has several important boundaries. First, the positive dataset is predominantly composed of animal toxins. Consequently, the reported performance is most directly applicable to this domain, whereas the generalizability of the approach to microbial and plant toxins remains to be established.

Second, the external validation included only 16 toxin sequences from a single species. Performance estimates from a dataset of this size are sensitive to individual predictions, and a single-species evaluation cannot fully characterize model behavior across diverse venomous taxa. Thus, although the *P. rubripinnis* results demonstrate generalization to an unseen proteome, they do not establish consistent performance across phylogenetically distant groups. Third, computational alanine scanning was applied to only one representative protein pair, so the highlighted positions should be regarded as model-derived hypotheses rather than experimentally validated determinants of toxicity. These limitations suggest several directions for future work. Evaluating NeoToxPred against a broader panel of venom proteomes would extend the present cross-species validation into a more systematic assessment of taxonomic transferability and determine whether the performance observed for *P. rubripinnis* is maintained across proteomes with distinct toxin repertoires. The candidate residues identified by alanine scanning could likewise be evaluated through targeted mutagenesis or, on a larger scale, deep mutational scanning. The latter approach could assess combinations of substitutions and thereby probe epistatic interactions that cannot be resolved by single-position perturbations. Finally, as curated datasets for microbial and plant toxin lineages become available, expanding the training data will help determine whether the negative-set design principles established here generalize beyond animal venoms. Despite these limitations, the findings support the central premise of this study. NeoToxPred can distinguish toxins from closely related non-toxic homologues using primary sequence information alone, without requiring the computationally intensive structural predictions used by some alternative methods. The comparative results further indicate that training-data construction, particularly the treatment of evolutionary relationships and sequence-length distributions, can be an important determinant of classifier performance. Therefore, careful curation of negative datasets should be considered alongside algorithmic architecture in the development and benchmarking of future toxin classifiers.

## Acknowledgments

This research was supported by the Korea Institute of Marine Science & Technology Promotion (KIMST), funded by the Ministry of Oceans and Fisheries (RS-2025-02292973); the National Research Foundation of Korea (NRF), funded by the Ministry of Science and ICT (RS-2025-02217886 and RS-2025-02309093); and the Bio & Medical Technology Development Program of the National Research Foundation (NRF), funded by the Korean government (MSIT) (No. RS-2025-24523548); and Global-Learning & Academic research institution for Master’s · PhD students, and Postdocs (LAMP) Program of the National Research Foundation of Korea (NRF) grant funded by the Ministry of Education (No. RS-2024-00442775)

## Author contributions

S.K. designed the study, conducted all computational experiments, and analyzed the data. S.K. wrote the original draft of the manuscript. C.P. and M.S.K. supervised the research, coordinated project execution, and critically revised the manuscript for intellectual content. All authors read and approved the final manuscript.

## Data availability

The source code and processed datasets required to reproduce and evaluate the findings of this study are openly available in the NeoToxPred GitHub repository at https://github.com/CSB-hub/NeoToxPred.

