## Supplemental Figs. Tables for "NeoToxPred: a fine-tuned protein language model with orthologous and length-stratified negative controls for robust toxicity classification"

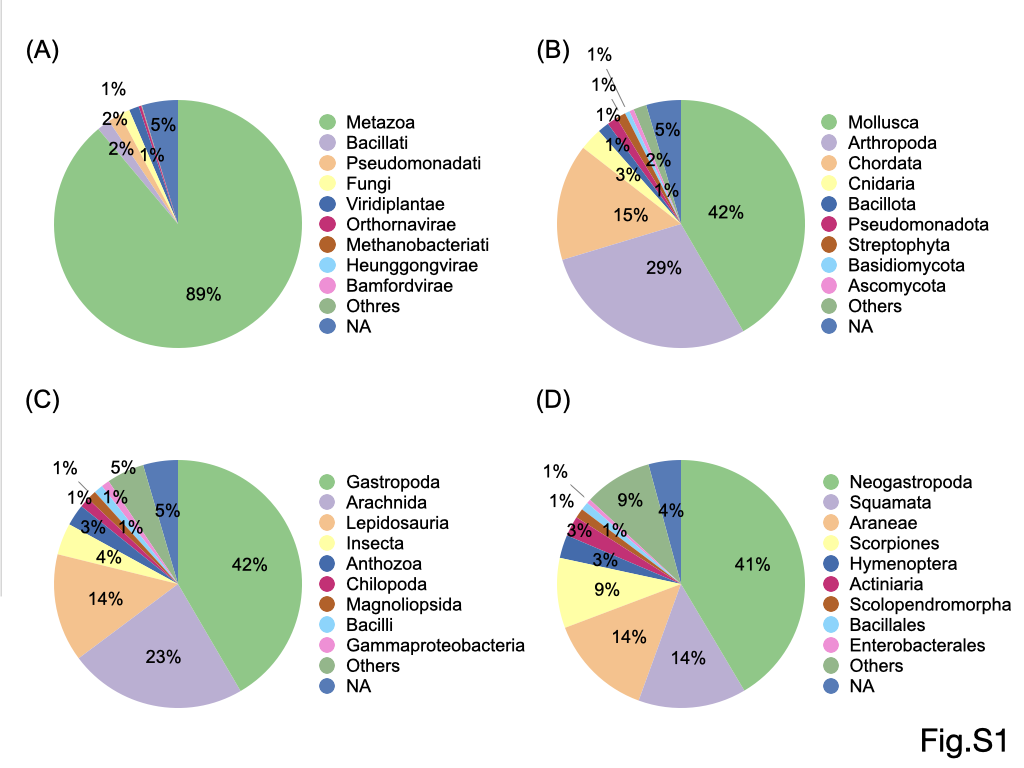

**Figure S1. Taxonomic distribution of the initial positive toxin dataset**. Pie charts delineate the distribution of the 16,687 non-redundant toxin sequences across four hierarchical taxonomic levels: (A) kingdom, (B) phylum, (C) class, and (D) order.

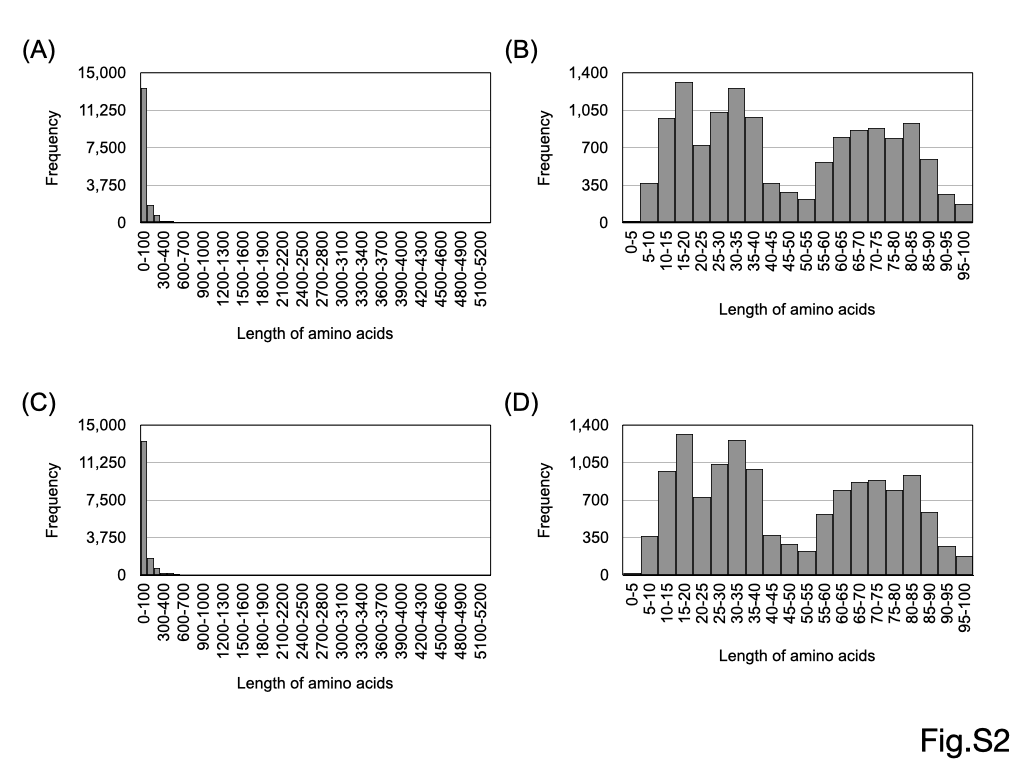

**Figure S2. Comparative sequence length distributions of the positive toxin and random negative datasets.** Histograms delineate sequence length frequencies for (A, B) the 16,687 non-redundant toxin sequences and (C, D) the 14,189 length-matched random negative sequences. Distributions are plotted across (A, C) the complete range of 0–5,200 residues and (B, D) a truncated range of 0–100 residues to highlight the short-peptide subset.

**Table S1. Number of sequences assigned to each data split.**

| Training set | Validation set | Test set |
| --- | --- | --- |
| 40,608 | 5,073 | 5,082 |

**Table S2. Performance of NeoToxPred across orthologous negative sets at four taxonomic levels.**

| Taxonomy level (n)^a^ | MCC | F1-score | Precision | Recall | Specificity | NPV |
| --- | --- | --- | --- | --- | --- | --- |
| Kingdom (826) | 0.838 (±0.046)^b^ | 0.915 (±0.034) | 0.910 (±0.066) | 0.921 (±0.021) | 0.923 (±0.042) | 0.921 (±0.036) |
| Phylum (5,137) | 0.820 (±0.046) | 0.906 (±0.024) | 0.909 (±0.051) | 0.904 (±0.024) | 0.913 (±0.056) | 0.913 (±0.008) |
| Class (10,188) | 0.824 (±0.030) | 0.915 (±0.018) | 0.923 (±0.041) | 0.907 (±0.019) | 0.922 (±0.041) | 0.897 (±0.051) |
| Order (20,171) | **0.873 (±0.013)** | **0.934 (±0.012)** | **0.942 (±0.016)** | **0.926 (±0.024)** | **0.946 (±0.013)** | **0.932 (±0.016)** |

^a^ Values in parentheses indicate the number of sequences in each orthologous negative set.

^b^ Values represent the mean ± standard deviation across five replicates.

Bold indicates the highest value for each metric across the four taxonomic levels.
